# Finding and exploring reproducible cell phenotypes with the Stemformatics data portal

**DOI:** 10.1101/2023.06.05.543705

**Authors:** Jarny Choi, Suzanne Butcher, Paul Angel, Jack Bransfield, Jake Barry, Noel Faux, Bobbie Shaban, Priyanka Pillai, Aleks Michalewicz, Christine Wells

## Abstract

Stemformatics is an established online data portal which hosts hundreds of curated gene expression datasets. It has been serving the stem cell research community for over a decade, by hosting transcriptional profiles of pluripotent and adult stem cells and their progeny from multiple tissues and derivation methods. The portal provides easy-to-use online tools to explore gene expression patterns in published data. In recent years, Stemformatics has shifted its focus from curation to collation and integration of public data with shared phenotypes. It now hosts several integrated expression atlases based on human myeloid cells, which allow for easy cross-dataset comparisons and discovery of emerging cell subsets and activation properties. The inclusion of laboratory-derived cell types enables users to benchmark their own data, to assist with cell-type standardisation or improve cell-derivation methods. The sample annotations have been greatly improved to enable better data integration, and the website has also undergone a major upgrade to modernise its visualisation tools. An application programming interface server also provides the data directly for computational users. Stemformatics is an open-source project and readily available at stemformatics.org.

## Introduction

Stemformatics is a gene expression data portal which hosts manually selected datasets of relevance to the international stem cell research community. The curation workflow includes careful re-annotation of experimental and sample metadata, using suitable standards where they exist. This in turn facilitates cross-dataset comparisons. Stemformatics reprocesses all hosted data from the source files, to standardise the data formats and assess uniform quality control metrics. Approximately 30% of public datasets reviewed by Stemformatics failed reprocessing or reannotation, because of ambiguities in the sample tables or because of poor quality primary data [1]. All these properties mean Stemformatics has been built on the FAIR (Findable, Accessible, Interoperable, and Reusable) guiding principles [2], and provides a unique resource to the stem cell research community. This is a key procedure to the FAIR data principles by which Stemformatics operates: not only reusing public data but adding domain knowledge and value to those published studies. Stemformatics is the only resource of its kind where induced pluripotent stem cell (iPSC), in vitro and in vivo derived blood cells can be explored and compared together at the transcriptional level [1][2] (Table 1).

**Table 1.**
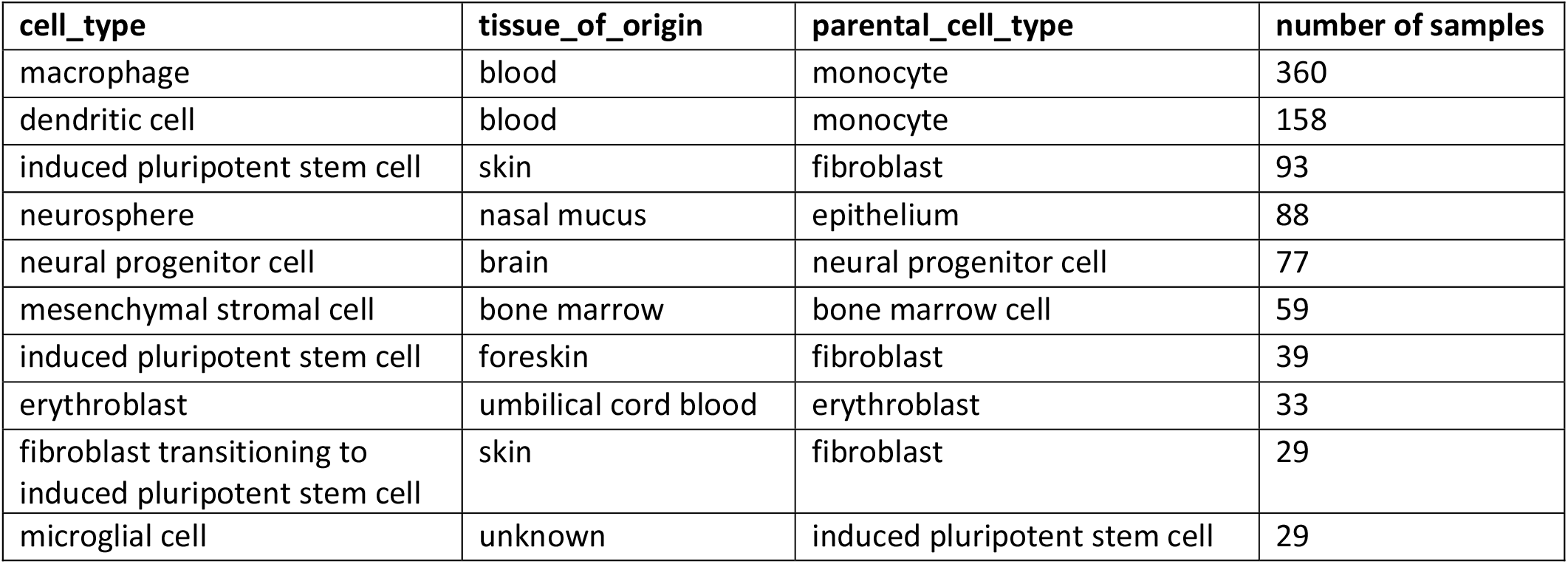
Top 10 cell types in Stemformatics ordered by number of samples, where each cell type is subset using tissue of origin and parental cell type (applicable when the cell has been differentiated from this parent). Improving annotation is an ongoing effort, and these numbers will vary as further samples are annotated.

| cell_type | tissue_of_origin | parental_cell_type | number of samples |
| --- | --- | --- | --- |
| macrophage | blood | monocyte | 360 |
| dendritic cell | blood | monocyte | 158 |
| induced pluripotent stem cell | skin | fibroblast | 93 |
| neurosphere | nasal mucus | epithelium | 88 |
| neural progenitor cell | brain | neural progenitor cell | 77 |
| mesenchymal stromal cell | bone marrow | bone marrow cell | 59 |
| induced pluripotent stem cell | foreskin | fibroblast | 39 |
| erythroblast | umbilical cord blood | erythroblast | 33 |
| fibroblast transitioning to induced pluripotent stem cell | skin | fibroblast | 29 |
| microglial cell | unknown | induced pluripotent stem cell | 29 |

Stemformatics was first introduced to the stem cell community in 2011 as the collaboration platform for Stem Cells Australia. It hosted the Project Grandiose stem cell reprogramming consortium in 2014 [3][4] and was updated in 2019 to allow for cross-dataset comparisons at the level of an individual gene. The focus of the site has since moved from collection and curation of relevant datasets to include integration of data into cohesive atlases. The novel integrated atlases facilitate comparisons between samples from different datasets to give biological insight into shared phenotypes, such as genes that mark shared cell identities or differentiation states. To support this integration, we have assigned uniform nomenclature to all sample metadata imported into the resource (Tables 1 and 2). User interviews were also conducted to improve functionality and usability of the website.

**Table 2.** Example changes in sample annotation from the previous version where uniform nomenclature has been applied. Where the terminology contained cell of origin for differentiated cell, such as “hESC-derived neuron”, we’ve added this to the “parental_cell_type” field.

| Term before update | After update |
| --- | --- |
| CD4 T cell | CD4-positive T lymphocyte |
| B cell | B-lymphocyte |
| Blood dendritic cells | dendritic cell |
| human embryonic stem cell | embryonic stem cell |
| Embryonic stem cell | embryonic stem cell |
| hESC-derived neuron | neuron (+ parental_cell_type = embryonic stem cell) |
| primary human neural progenitor | neural progenitor cell |
| BMDM | macrophage (+ tissue_of_origin = bone marrow) |
| primary skin fibroblast | fibroblast (+ tissue_of_origin = skin) |
| hematopoietic multipotent progenitor | made more specific or changed to hematopoietic precursor cell (or, if FACS sorted, CD34-positive precursor cell) |

We manually curated sample metadata to harmonise data groupings, expose experimental validation of cell type, and improve user experiences when searching for specific cell types (Table 2). To leverage existing knowledge in this area, we used the Ontology Lookup Service tool [5] from (European Molecular Biology Laboratory-European Bioinformatics Institute (EMBL-EBI). For disease state, we sourced annotations from the Human Disease Ontology [7]. For tissue of origin, we sourced annotations from the Brenda Tissue Ontology [8]. For cell type, parental cell type and final cell type, we used the Cell Ontology [9]. Where known cell lines were included, we used Cellosaurus as an identifier reference [6]. This enables deep dives into specific systems such as iPSC derived myeloid cells and allows for more robust sample search functionalities on the website.

The website has been completely redesigned using modern infrastructure and technologies. It now runs a separate API (application programming interface) server, which can be used to query the data directly for advanced users. The user interface server uses modern JavaScript technologies for easier development and maintenance. The system and the code have also been designed to work as a base for other data portals, suitable for research-oriented environments. This report provides an overview of the user interfaces on the site, and examples of the biology available for exploration.

## Exploring the data

Stemformatics website invites users to explore the data using visual tools in addition to the traditional text-based searches (Fig 1). The Visual Data Explorer (/datasets/explore, Fig 1B) is an interactive sunburst plot which can represent hierarchical relationships with its concentric circles, such as parental cell types which give rise to final cell types in a differentiation process. Clicking on any of the elements shows how many datasets and samples correspond to that value and the user can then readily explore these datasets.

**Fig 1.**
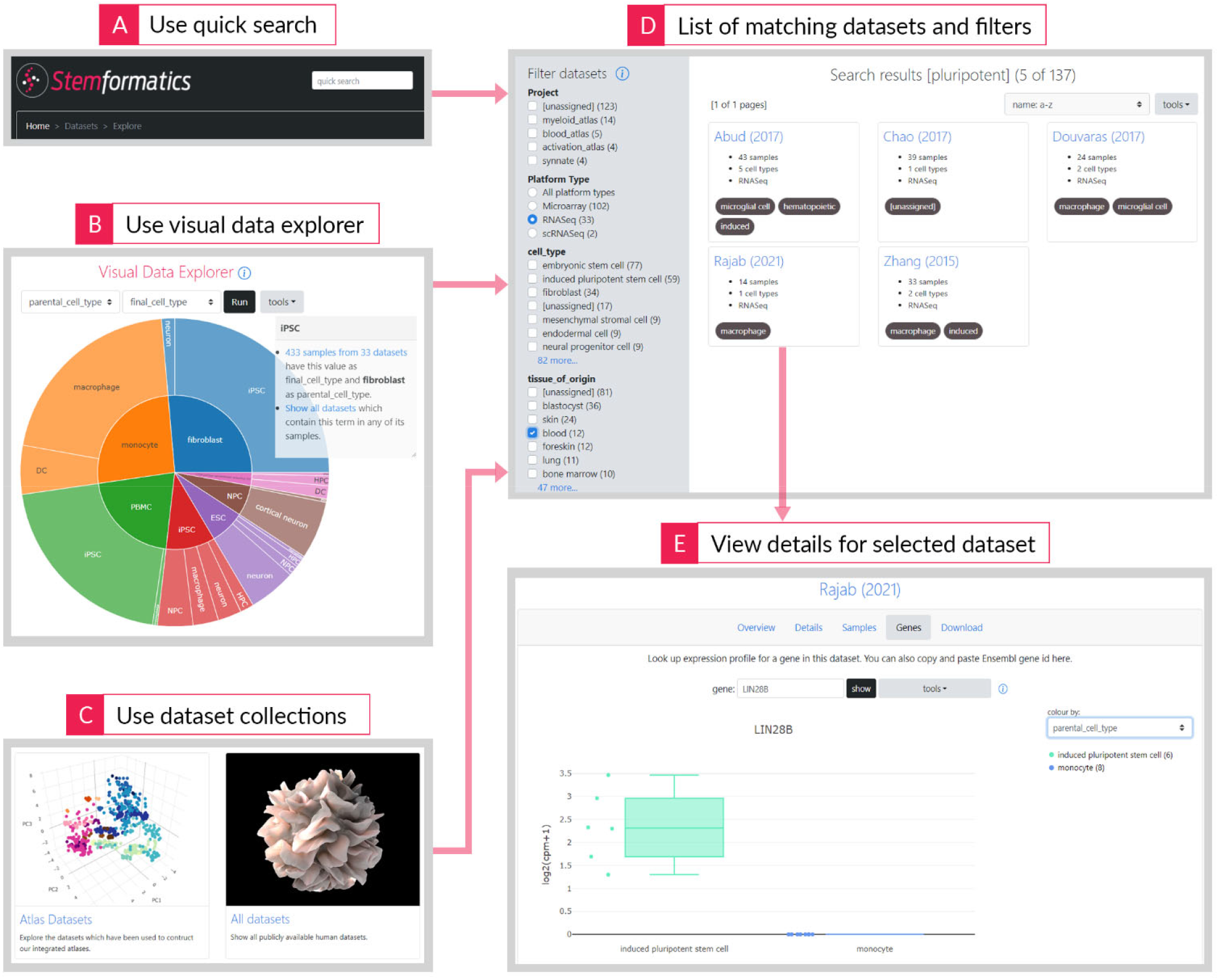
Exploring datasets hosted on Stemformatics can happen from multiple starting points. Quick search allows for word-based search through dataset metadata (A). Visual data explorer allows for an interactive view of related samples (B). And collections allow access to datasets grouped together under a project (C). The list of datasets returned can then be further filtered – here we used the term “pluripotent” in the quick search, then narrowed down the results to RNA-seq data from the blood (D), and clicking on the individual dataset allows for PCA and gene expression plots, metadata views and data downloads (E). In this example, the LIN28B gene shows differential expression between iPSC derived and in vivo derived macrophages.

Collections (/datasets/collections, Fig 1C) are groups of datasets gathered for some commonality, such as containing leukemia cells, or used in building the integrated atlases. Each dataset can be assigned multiple project tags, which can also be used to group datasets together. This feature also serves collaborative projects.

Once the user selects a collection of datasets to view, they can be further filtered (/datasets/filter, Fig 1D) to a smaller subset based on properties of interest, such as particular sequencing technologies or cell types. Finally clicking on the individual dataset from this filter page shows details on the dataset, which includes the principal components analysis (PCA) plot of the dataset to visualise the sample relationships to each other.

Other tools on the dataset include the gene expression plot, which shows the expression profile of any gene in the dataset, finding correlated genes (Fig 1E), and viewing the dataset and sample metadata. The expression data and sample metadata can be downloaded as text files.

To explore genes based on expression profiles, two types of searches are available: show highly expressed genes starting from a sample group (/genes/sampletogenes)or start from a gene to find samples where it is highly expressed (/genes/genetosamples). We wanted to leverage the large number of datasets in Stemformatics to show patterns which are repeatedly seen across multiple datasets. Hence the output from these searches is shown in a hierarchical manner: when a gene is found to be highly expressed in macrophages for example, multiple datasets are shown under macrophages for that gene.

## Integrated atlases

We were motivated to build an integrated atlas of myeloid cells in order to compare iPSC derived myeloid cells to their in vitro and in vivo counterparts. We wanted to explore how different sample sources or derivation methods may have an impact on the cell identity, and this was not possible when many of these features of interest were scattered all over multiple datasets. The old Stemformatics portal hosted multiple datasets in parallel, where the user could access each dataset separately. Comparing data across datasets would bring valuable biological insight, but technical differences in the measurements and scale of data prevented direct comparisons.

Many different methods are available for batch correcting multiple bulk transcriptome datasets to integrate them. We chose variance partition [10], applied after rank transforming each expression matrix. This method produced robust integrated atlases with real biological clusters [11]. We were then able to observe emerging properties from the combined data, such as finding that cord-blood derived dendritic cells retained an in vitro identity, likely from lack of appropriate growth factor signalling [12].

Leveraging the scalability of this approach, we created three integrated atlases: Blood, Myeloid and Dendritic Cell, and these are hosted on Stemformatics website (/atlas/blood, /atlas/myeloid, /atlas/dc). Each atlas comes with a set of additional annotations relevant for that system, and the web page is full of features for easy exploration and usage (Fig 2). The PCA plot of the atlas is the default view and can be used to visualise the relationship of cells to each other. Cells can be coloured based on predefined categories, such as cell type, tissue of origin or parental cell source (Fig 2A). The data can be explored further by customising the sample groups shown on the graph. This feature allows users to combine categories – for example cell type and tissue source – so as to group and view samples annotated with these terms (Fig 2B). This feature allows the user to explore more precise relationships between cellular behaviours and experimental factors, including discovery of the influence of various experimental factors on a cell’s transcriptional state. Because these states can be visualised across many independent datasets, the reproducibility of these behaviours are straight forward to assess. The user can also search for individual genes in the atlas and view its expression as a box or violin plot (Fig 2D) or as a colour gradient imposed on the PCA plot (Fig 2E). For example, Fig 2D shows that the cDC1 marker XCR1 has variable expression between blood (high expression) or bone marrow (low expression). This may assist prospective isolation of these cells from tissues where commonly used markers are not suitable.

**Fig 2.**
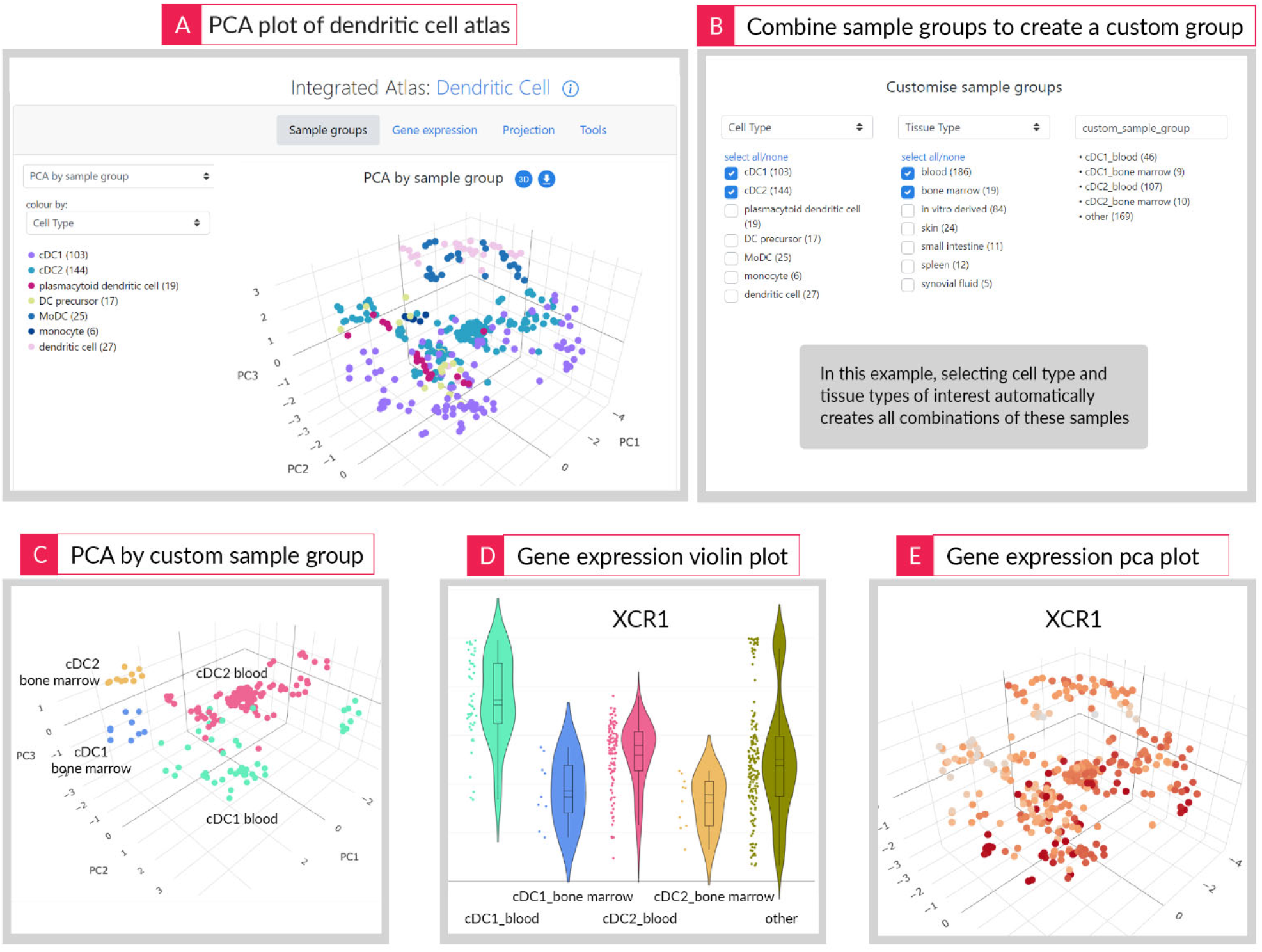
Different visualisations options available on a Stemformatics atlas page. PCA plot (A) is the default view which allows for cell type and cell state comparisons. In this example, we investigate tissue specificity of cDC1 and cDC2 by creating a customised sample group which combines cell type and tissue type (B). This leads us to view that samples cluster by tissue type within each cell type (C). Querying the expression of XCR1 gene shows higher expression in blood than bone marrow within each of these cell types (D,E).

Transparency is core to the Stemformatics user experience, so when users find a sample of interest on the atlas, a simple double click on any cell in the PCA plot will show the origin of that cell – which dataset it came from and all its annotations. The user can also look at the full list of the datasets which were used to construct an atlas. Likewise, all of the data used to construct each atlas can be downloaded as text files, including the exact colours used to render the plots.

## Benchmarking data using integrated atlases

As new gene expression datasets are produced, it would be very insightful to be able to compare their transcriptional profiles to an established set of references. Stemformatics integrated atlases provide a unique resource that can act as such references. Often a reference dataset is a large study from a single laboratory, which means it may represent a single platform or tissue of origin for example. Hence when a query is made against such a reference, these variables may act as confounders. In contrast, a Stemformatics atlas is built from a manually curated list of many datasets and the same biological samples are represented across different platforms and other experimental factors.

Stemformatics website makes it easy to benchmark a bulk gene expression dataset against an atlas, where the expression matrix and sample table can be uploaded as tab separated text files. Two independent methods are used simultaneously on each query:

1. Query samples are projected onto the PCA space of the atlas and results are shown in the PCA plot. Fig 3 shows an example where data from Monkley et.al. [13] have been projected onto the Stemformatics Dendritic Cell Atlas. Monkely et.al. derived macrophages and dendritic cells.) from induced pluripotent stem cells (iPSCs) and obtained their molecular profiles through bulk RNA-seq. The projection shows their iPSC derived cells remain transcriptionally close to other in vitro derived macrophages in the atlas, whereas their peripheral blood mononuclear cells (PBMCs) are close to the in vivo sample sources in the atlas. Interestingly, even their PMBC derived cells assume the in vitro identity after being cultured for several days. This observation is however consistent with other cultured cells which were used to construct the atlas – that culturing can shift a cell’s identity significantly away from their direct in vivo counterparts.

**Fig 3.**
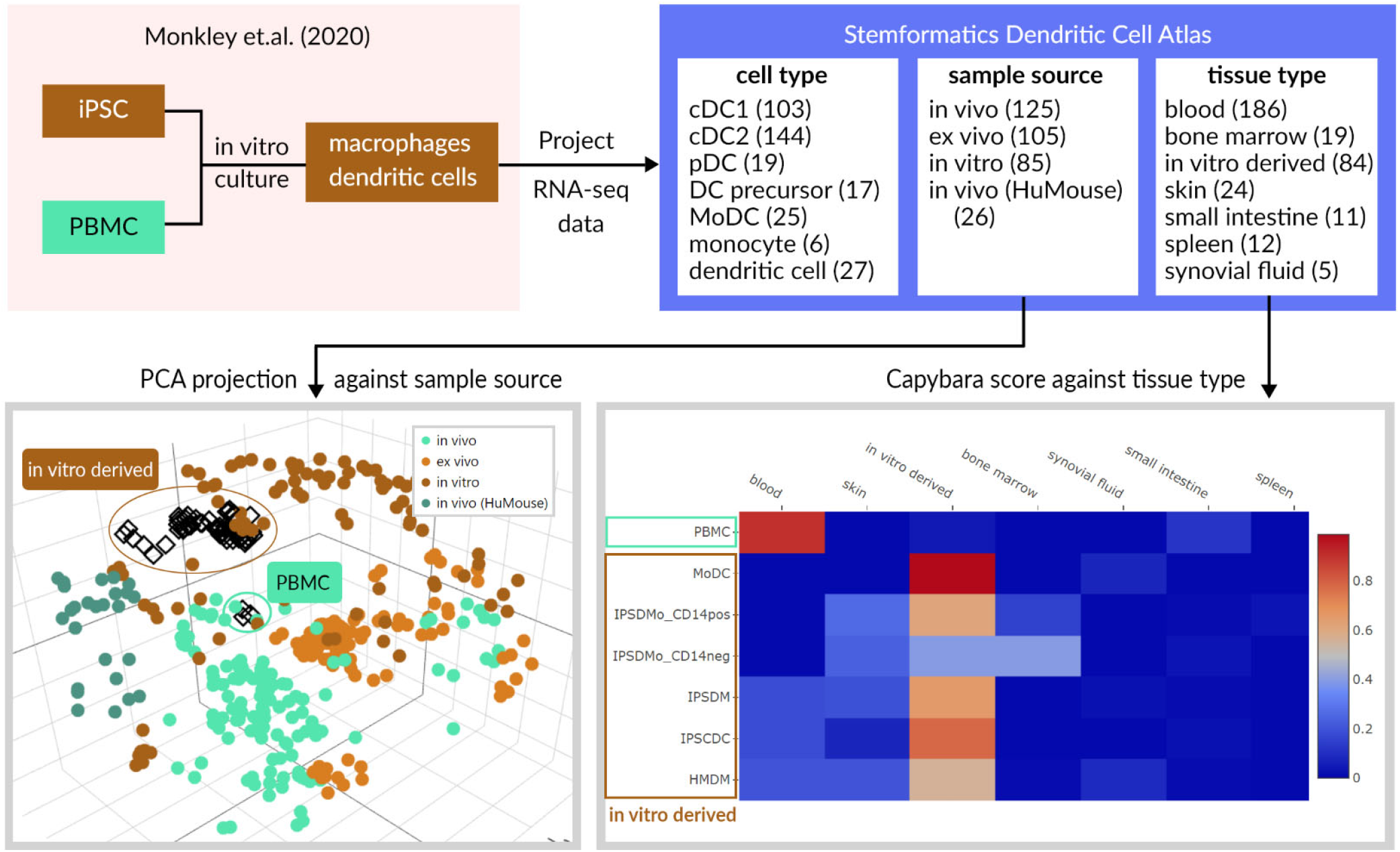
Results of projecting an external data onto the Stemformatics atlas demonstrate its utility as a benchmarking tool. Monkley et.al. is a dataset containing iPSC and in vivo derived myeloid cells, which has been projected onto the Stemformatics Dendritic Cell Atlas. Two independent methods of classifications are applied simultaneously on the website, leading to the visualisation in PCA space of projected cells (bottom left - projected cells are diamond shapes) and heatmap of Capybara scores (bottom right). The user can dynamically change the reference sample group after the projection has been made, to easily compare their samples against various cell phenotypes in the atlas. Since each projection is independent of other projections, this method avoids any batch effect that may result from integrating the query data with reference. It also provides an intuitive way to understand and visualise the comparison in terms of the principal components of the reference.
2. Capybara [14] is used to produce a score (between 0 and 1) for each query sample against the reference samples. Capybara uses constrained linear regression to produce the scores as a continuous variable and is particularly well suited for this analysis. We implemented a python version of its key function for use against Stemformatics atlas as a reference. The scores are also calculated against all sample groups in the atlas, not just cell type, hence concordance with tissue type or sample source (in vitro, ex vivo, in vitro) are also shown. Fig 3 shows the Capybara result for querying the Monkley dataset against the tissue types in the Stemformatics Dendric Cell atlas. It shows high concordance between their in vitro derived cells and the “in vitro derived” tissue type in the atlas.

Single cell RNA-seq data can also be queried against the atlas. To make the range of expression values more compatible, single cell data should be either pseudo-bulk aggregated or imputed first. Deng et.al. [15] has detailed description of how this could be performed and the Stemformatics website also provides a vignette. Here we projected a subset of the Rosa et.al. [16] data onto the Stemformatics Myeloid Atlas after pseudo-bulk aggregation (Fig S1). Rosa et.al. cultured DCs from human embryonic fibroblasts (HEF) and profiled them using single cell RNA-seq. The PCA projection of their HEF derived DCs onto the Stemformatics myeloid atlas shows a correlation between the age of the cells and the “activation axis” of the atlas which correspond to maturity of cell types. These cells also project closely with the in vitro derived myeloid cells in the atlas, in contrast to their peripheral blood derived controls, which project closely with the vivo derived dendritic cells in the atlas.

Rosa et.al. used Villani et.al. [17] data to benchmark their cells in their paper. We propose that Stemformatics atlases would also provide excellent references in this case, since they contain iPSC derived cells that Villani et.al. does not. Furthermore, the Stemformatics atlases employ multiple comparison methods and easy-to-interpret visualisations from simple point and click tools. Note that all data for each atlas are available for download as text files, so custom methods of benchmarking can easily be applied to these references.

## API Access and System Design

Stemformatics is built as two separate applications: an application programming interface (API) server which hosts all the data, and a user interface (UI) server which hosts the website (Fig 4). This is a common design paradigm for data heavy websites, and allows for flexibility in both development and maintenance of the system.

**Fig 4.**
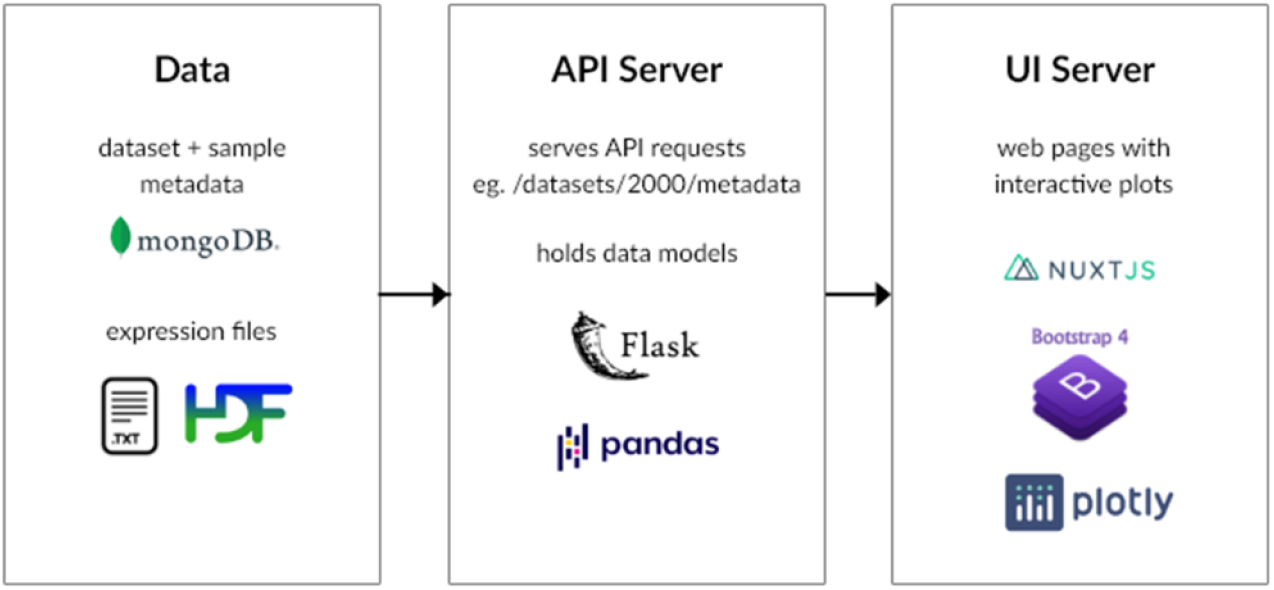
A schematic of the Stemformatics infrastructure, which separates the API server from the UI server and is built on modern full stack technologies, such as Flask, MongoDB, Bootstrap and Nuxt.

The API server provides direct access to the data without going through the website, making it convenient for computational biologists and bioinformaticians to access the data and perform analyses. An example use-case may be to search for all RNA-seq datasets containing a particular cell type and download the corresponding counts per million (CPM) matrices for downstream analyses. In R, this code may look like this (note: each dataset in Stemformatics has a unique 4-digit number as an identifier):

~~~
# R example
library(httr)
library(jsonlite)
response = GET(“https://api.stemformatics.org/search/samples?query_string=microglia&field=cell_type,dataset_id”)
data = content(response) # data is a list
datasetIds = unique(sapply(data, function(x) getElement(x,’dataset_id’)))
for (datasetId in datasetIds) {
 df = read.csv(paste0(“https://api-dev.stemformatics.org/datasets/“, datasetId, “/expression?as_file=true”),
 sep=‘\t’, row.names=1)
 write.table(cbind(id=rownames(df),df), file = paste0(datasetId, “.tsv”), sep=“\t”, quote=F, row.names=F)
}
~~~

The API server is built on Flask-restful [18]. It uses pandas [19] extensively to manipulate data frames. The dataset and sample metadata are held as collections in a MongoDB [20], while expression files are stored as text and hdf5 [21] files. The UI server is built on Nuxt JS [22], using BootstrapVue [23] to build components easily. Most of the plots are performed by Plotly [24]. These tools are designed for modern browsers and user interfaces.

Its open-source code base makes an excellent reference for similar efforts and is available at github.com/wellslab/s4m-api and github.com/wellslab/s4m-ui. Full stack development within a research environment comes with some challenges, which include small (or quite often one person) teams, rapid turnover of personnel and resource issues for continued maintenance. The Stemformatics code has been designed to address some of these issues, based on a decade of experience in maintaining resources like this. The key principle is to strike a good balance between under or over-engineering the system. Under-engineering creates bloated code with highly dependent variables and states which are difficult to change and maintain. This can be avoided by breaking up complex pages or classes into independent components. Over-engineering creates code only understandable with highly specialised skills, and can be avoided by greater transparency and less layers of abstraction.

Stemformatics also supports private datasets. These are datasets flagged as private in the dataset metadata, which are only accessible through an account login. We use this feature to share data with collaborators prior to publication.

## Discussion

FAIR data principles suggest that data should not only be findable, but also reusable. The Stemformatics platform provides an example of data reuse for community benefit, where we leverage data from otherwise obsolete technology platforms to build new insights into the shared behaviour of cells across different laboratories, derivation methods and activation states. The ethos of the Stemformatics team is to provide a community-facing platform that is easy for biological experts to use without requiring complex bioinformatics software. All of the infrastructure, data processing methods and primary data sources are provided, and APIs facilitate access from external computational workflows.

By hosting many independent datasets with varied and overlapping covariates and annotating these in a consistent manner, Stemformatics provides an alternative scale of big data to single cell RNA-seq studies. With Stemformatics, gene networks which separate the samples are often reflecting the nuanced cell states based on the multiple covariates, such as the tissue source or activation status of the cell. Macrophages may be differentiated in vitro from different tissues, under different stimuli and disease conditions, for example. Observing these patterns repeatedly across many datasets increases confidence in their biological signal.

As more single cell RNA-seq datasets are created, the relevance of bulk RNA-seq and microarray studies should be evaluated within the appropriate context. In the stem cell research field, in vitro models are extensively used, and these models have been developed and refined for over a decade. Hence the data captured by older technologies hold many experimental factors which influence each model. These factors may be viewed as covariates if we are making inferences on the output of the models. It may be expected that single cell datasets will also cover these covariates in future (see Alsinet et.al. [25] for example), but currently the bulk studies far outnumber single cell studies, and their cheaper cost means they will continue to be produced. The former are more suitable for discovering heterogeneity within the models and the gene networks will often reflect the cell type differences.

No single tool or reference is likely to capture the precise states of many cells, hence it is important to understand their limitations. We recommend that multiple tools and references should be used to cross-check the results for cell type classification, rather than relying on a single source. Stemformatics atlases provide an important resource which leverages the wealth of built-up knowledge in the community. Having high levels of manual curation means Stemformatics atlases focus more deeply on particular models of interest by design (only myeloid cells for example). This should be taken into account when using Stemformatics atlases for benchmarking.

## Conclusion and future directions

Stemformatics is a unique resource which empowers the stem cell and immunology research communities by hosting high quality datasets and easy to use online tools. It also provides integrated expression atlases focused on myeloid cells derived from multiple sources and conditions. This makes it possible to compare iPSC derived macrophages to their in vivo counterparts, for example. These atlases also serve as excellent benchmarking tools for related cells.

We plan to build on the integrated atlases to uncover key regulators of cell states. Since each atlas is based on deep curation of cellular phenotypes, it can expose both the important genes for cell states as well as the experimental factors which define these.

Recently updated website has been built on modern full stack technologies and the API server allows for programmatic access to the data. Stemformatics has been built on FAIR principles, and all its data and code are easily accessible and can serve as a template for similar projects.

## Acknowledgements

We would like to acknowledge other members of the Synnate team at the Hudson Institute for providing input, especially Jamie Gearing.

## Author contributions

JC developed the methods, database and software and wrote the manuscript. SB annotated and processed data. PA developed methods. JB and JB developed software. NF, AM and PP developed annotation guidelines. BS developed annotation guidelines and visualisation prototypes. CW conceived the project, annotated data, funded the project and wrote the manuscript.

