## Supplementary Materials for "Finding and exploring reproducible cell phenotypes with the Stemformatics data portal"

**Fig S1.** Single cell RNA-seq data can also be queried against a Stemformatics atlas for benchmarking. Rosa et.al. is a single cell RNA-seq dataset containing dendritic cells which have been reprogrammed from human embryonic fibroblasts (together with fluorescence activated cell sorting (FACS) sorted cells from peripheral blood as controls). When we projected these data onto the Stemformatics Myeloid Atlas after pseudo-bulk aggregation, we found the reprogrammed dendritic cells clustering with in vitro derived myeloid cells in the atlas. In contrast, the peripheral blood derived cells cluster with other in vivo derived dendritic cells in the atlas. Since the first principal component of this atlas separates progenitors from mature cells, it is interesting to note that this direction correlates well with day of extraction for the reprogrammed cells.

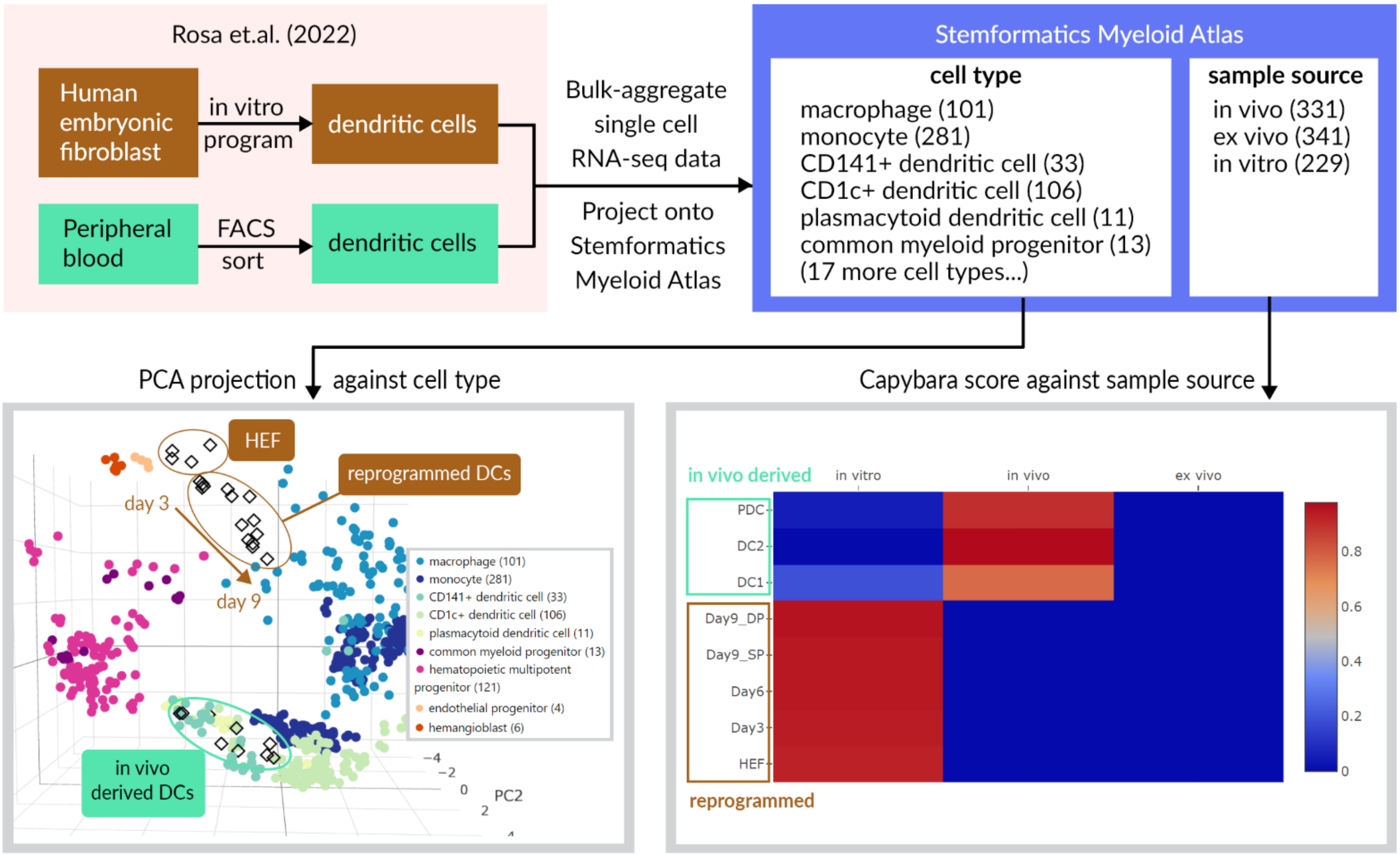
